# Fingertip–Surface Interfacial Shear Stress Varies with Sliding Conditions and Electrostatic Actuation

**DOI:** 10.64898/2026.08.13.744590

**Authors:** Celal Umut Kenanoglu, Yasemin Vardar

## Abstract

Fingertip friction plays a central role in tactile exploration and object manipulation. During sliding, tangential force depends jointly on the real contact area and the interfacial shear stress, both of which can be influenced by sliding conditions. However, changes in fingertip friction are often interpreted primarily through changes in real contact area, whereas the accompanying changes in interfacial shear stress remain less well characterized. This gap is especially relevant for electrostatic surface haptic displays, which modulate fingertip friction by applying a voltage between the finger and the touch surface. Here, we experimentally quantify the mean interfacial shear stress of a sliding fingertip on an electrostatically actuated touchscreen using simultaneous measurements of tangential force and optically resolved real contact area. Ten participants performed sliding trials across three speeds and three normal forces with and without electrostatic actuation. Interfacial shear stress increased with speed and decreased with normal force; in both cases, these trends arose because real contact area varied more strongly than tangential force. Electrostatic actuation further reduced interfacial shear stress, as increasing voltage produced a larger increase in real contact area than in tangential force. These findings show that interfacial shear stress varies systematically with sliding conditions and electrostatic actuation, clarifying how changes in real contact area and interfacial shear stress combine to shape fingertip–surface friction.

## I. Introduction

**F**RICTION between the fingertip and a contact surface plays a central role in precisely manipulating objects and in perceiving tactile properties of surfaces [1]–[3], and has therefore attracted attention in robotics, haptics, and tribology [1], [4], [5]. Despite an extensive body of research, interpreting the measured tangential force remains challenging because it emerges from the mechanics of soft finger contact. As the fingerpad is compliant and viscoelastic, sliding conditions (e.g., applied normal force and sliding speed) can alter contact formation, load distribution, and the resulting frictional response [3], [6]–[8].

A common way to interpret the tangential force is to express it via the Bowden and Tabor formula, as the product of the real contact area and the interfacial shear stress, *F*_*t*_ = *τ A*, where *A* denotes the real contact area formed by microscopic asperity junctions, and *τ* represents the interfacial shear stress during sliding [9]. Accordingly, the real contact area has received considerable attention as a key quantity in friction and contact mechanics [10]–[12]. However, this formulation gives real contact area and interfacial shear stress equal weight, meaning a change in tangential force can arise from a change in either quantity, or both. Previous works have demonstrated that the shear stress of soft contacts can vary with interaction conditions, including contact pressure in inner-forearm contacts [13], sliding velocity in rubber contacts [14], [15], dynamic loading frequency in fingertip contacts [16], and electrostatic actuation during fingertip sliding [17]. Therefore, interpreting changes in fingertip friction requires examining how real contact area and interfacial shear stress vary relative to one another.

Among these interaction conditions, normal force and sliding speed are the most extensively studied for fingertip contact. Previous studies have shown that fingertip friction and real contact area vary with normal force, sliding speed, skin condition, and surface properties [1], [3], [7], [12], [16]–[20]. Increasing normal force generally increases both tangential force and real contact area, although the friction coefficient often decreases with load. The effect of sliding speed is more condition-dependent: fingertip friction can exhibit non-monotonic behavior, and contact formation may be influenced by rate-dependent skin deformation, viscoelasticity, hydration, occlusion, and the direction of exploration [3], [7], [18]– [21]. Moreover, tangential force and real contact area do not necessarily change proportionally across force and speed conditions. Specifically, it remains unclear whether force- and speed-dependent changes in fingertip friction arise primarily from changes in real contact area, changes in mean interfacial shear stress, or their combined variation. This problem has not been systematically quantified during steady fingertip sliding on glass across controlled normal-force and sliding-speed conditions. In addition, inter-participant differences in skin properties, finger geometry, hydration, and contact behavior may further affect the relative contributions of real contact area and interfacial shear stress.

Beyond its dependence on normal force and sliding speed, fingertip tangential force can also be actively modulated using surface-haptic techniques. Electrostatic actuation is one of the most widely used approaches for this purpose; it modulates fingertip–surface friction by applying an alternating voltage to a conductive layer of the touchscreen [5], [22]. Previous studies have shown that electrostatic friction modulation depends on actuation voltage, frequency, sliding speed, and normal force [5], [17], [23]–[28]. The increase in tangential force under electrostatic actuation is commonly attributed to electrostatic attraction, which alters the fingertip contact mechanics and can increase the real contact area [11], [17], [25]. However, as tangential force depends jointly on real contact area and mean interfacial shear stress, changes in real contact area alone may not fully describe the interfacial response to electrostatic actuation. Our previous work showed that electrostatic actuation changed the mean interfacial shear stress when actuation frequency was varied at fixed sliding speed, normal force, and voltage [17]. Yet, it remains unclear how the change in *τ* under electrostatic actuation depends on sliding speed and normal force, and whether this change scales systematically with actuation voltage.

In this paper, we experimentally investigate how the mean interfacial shear stress of a sliding fingertip depends on sliding conditions and electrostatic actuation. We combine tangential force measurements with optically resolved real-contact-area measurements to calculate *τ* across three sliding speeds and three normal forces in ten participants. We first examine how *τ* varies under voltage-off conditions and relate these trends to the relative changes in tangential force and real contact area. We then test whether electrostatic actuation modifies this interfacial shear stress across the same speed–force conditions. Finally, a voltage-sweep experiment is used to examine how actuation amplitude changes the balance between tangential force, real contact area, and interfacial shear stress. Together, these analyses show that interfacial shear stress is not constant during fingertip sliding, but varies systematically with sliding conditions and electrostatic actuation.

## II. Materials and Methods

The dataset analyzed in this study was collected using the same setup and protocol (Fig. 1) as in our previous work [28]; here the key details are summarized for completeness. In addition to this experiment, a new voltage-sweep experiment was conducted as part of the present study. Both experiments were conducted in accordance with the Declaration of Helsinki and were approved by the Ethics Council of TU Delft (application no. 5108). All participants provided informed consent.

**Fig. 1:**
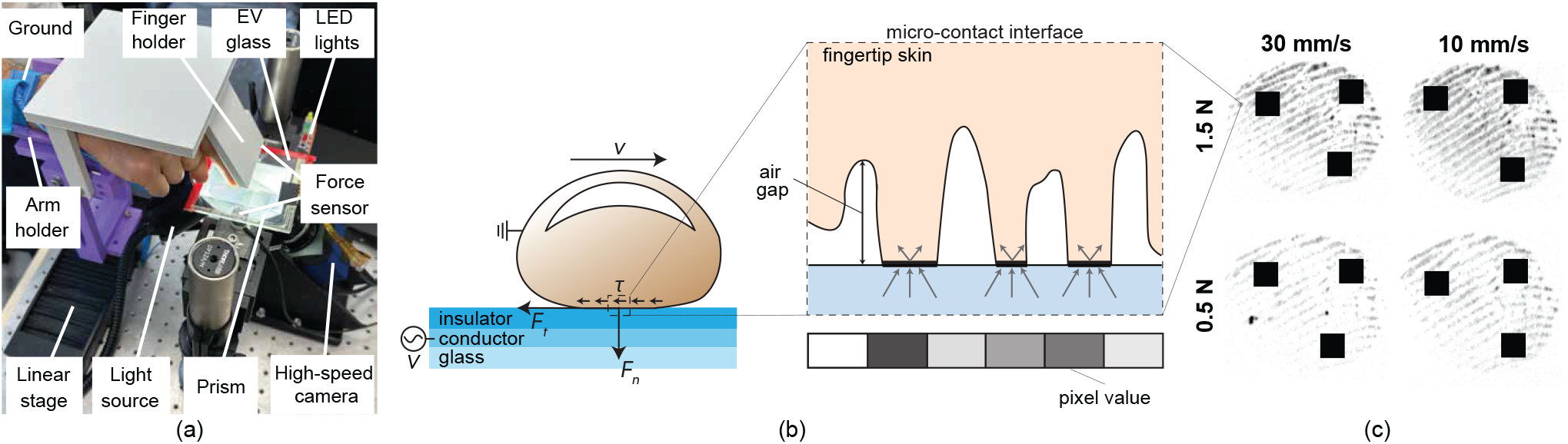
Experimental setup and force–area interpretation of fingertip sliding on an electrostatic touchscreen. (a) Experimental setup for simultaneous force and FTIR-based contact-area measurements. (b) Schematic of the fingertip–touchscreen interface. The fingertip slides with velocity *v* under applied normal force *F*_*n*_, while tangential force *F*_*t*_ is measured along the sliding direction. Interfacial shear stress *τ* describes the tangential load normalized by the measured real contact area. In the microcontact inset, the black regions indicate optically resolved intimate contacts, which define the optically resolved real contact area. Within the prism, light is totally internally reflected except at microscopic ridge asperities in direct contact with the prism surface. At these contact points, scattering disrupts total internal reflection, resulting in dark pixels, whereas non-contact regions appear bright. (c) Example finger images, where darker regions indicate greater real contact area.

### A. Experimental Setup

During the measurements, participants slid their right index finger across a capacitive touchscreen (SCT3250, 3M), whose conductive layer was excited by an alternating voltage. The excitation signal was generated using a data-acquisition board (SCB-68A, NI Inc.) at a sampling rate of 10 kHz and amplified using a high-voltage amplifier (9200A, Tabor Electronics). Participants wore an anti-static wrist strap. The touchscreen was rigidly mounted on two six-axis force/torque sensors (Nano17 Titanium, ATI Industrial Automation), which measured the normal and tangential interaction forces. Force signals were recorded using a separate data-acquisition card (PCIe-6321, NI Inc.) at 10 kHz. The finger motion was imposed by a motorized linear stage (NRT150/M, Thorlabs), which constrained the finger trajectory and maintained an angle of 60°. Data acquisition, stage control, and camera triggering were synchronized in MATLAB/Simulink. The real contact area was measured from below using frustrated total internal reflection (FTIR) imaging [16]. The glass plate was illuminated using an LED light source (KL 2500, Schott), and the contact images were recorded with a high-speed camera (MotionBLITZ EoSens mini2, Mikrotron) and lens (LM16HC, Kowa) at 1000 frames per second. The setup was mounted on damped posts on an optical breadboard to reduce external vibration effects.

### B. Experimental Protocol

The first experiment included ten participants (seven men and three women), with a mean age of 27, SD: ±2.45. Before testing, participants washed their hands and then dried their fingertips with a microfiber cloth. In each trial, the finger was moved laterally across the touchscreen at a constant speed of 10, 20, or 30 mm/s. Participants were instructed to maintain a target normal force of 0.5, 1.0, or 1.5 N using real-time visual feedback computed from the force-sensor signal. The feedback was provided using LEDs: yellow indicated forces more than 10% below the target, green indicated forces within the ±10% tolerance band, and red indicated forces more than 10% above the target. Each speed–force condition was repeated three times. Trials were retained only when the measured normal force remained within ±10% of the target and the fingerprint image was sufficiently visible for estimation.

To compare interfacial shear stress with and without electrostatic actuation, each sliding trial contained a voltage-off segment followed by a voltage-on segment, allowing direct comparison of the two actuation states within the same sliding trial and operating condition. The voltage-on segment used a 100 V_*peak*_, 75 Hz sinusoidal signal applied to the conductive layer of the touchscreen. To reduce moisture-related variability across trials, a small fan was used between trials to help maintain consistent skin dryness.

A separate voltage-sweep experiment was conducted to examine how actuation amplitude affected the force–area balance. This experiment involved one participant, aged 29 years, and used the same voltage-off–voltage-on trial structure as the first experiment. The actuation frequency was fixed at 75 Hz, while the peak voltage amplitude was varied across three levels: 50, 100, and 150 V. The sliding speed and target normal force were kept fixed at 20 mm/s and 1 N. Each voltage condition was repeated three times.

### C. Data Processing

A trigger was issued from Simulink when the finger entered the camera field of view and only the synchronized normal force (*F*_*n*_), tangential force (*F*_*t*_), and image data within this window were used for analysis. Before contact-area extraction, raw FTIR images were geometrically corrected. Radial lens distortion was removed using intrinsic parameters from checkerboard calibration, and a projective transformation was applied to obtain an equivalent top-down view [29]. The homography was calibrated by mapping the imaged ellipse of a circular rubber target placed on the glass to a true circle. The real contact area was then obtained from the corrected FTIR fingertip images following the procedure described in [12]. We use the term real contact area to denote the optically resolved contact regions at the interface; because contact area is scale-dependent, this estimate may differ from measurements obtained at smaller length scales [15], [30], and should therefore be considered as an optical approximation of the true contact area. This scale dependence may affect the absolute magnitude of real contact area; however, the present analysis focuses primarily on relative changes. The reported interfacial shear stress represents a real area-normalized tangential force (*τ* = |*F*_*t*_|*/A*), which captures condition-dependent changes in the measured force–area balance and is used here as a comparative measure across sliding and actuation conditions. For each trial, *F*_*t*_, *F*_*n*_, and *A* were computed as mean values over the corresponding analysis window. For voltage-on/voltage-off comparisons, 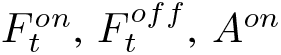, and *A*^*off*^ were computed as mean values over the corresponding voltage-on and voltage-off segments within the same sliding trial.

## III. Results and Discussion

The results show that interfacial shear stress varied systematically across the tested speed-force conditions (Fig. 2). Higher sliding speeds were associated with larger interfacial shear stress values, whereas increasing normal force generally reduced the interfacial shear stress, although this effect was comparatively smaller. These observations indicate that the effective mechanical parameters of the finger–surface interface depend on the sliding conditions rather than remaining constant. As interfacial shear stress reflects the combined evolution of tangential force and real contact area, its quantitative trend and mechanical basis are examined next.

**Fig. 2:**
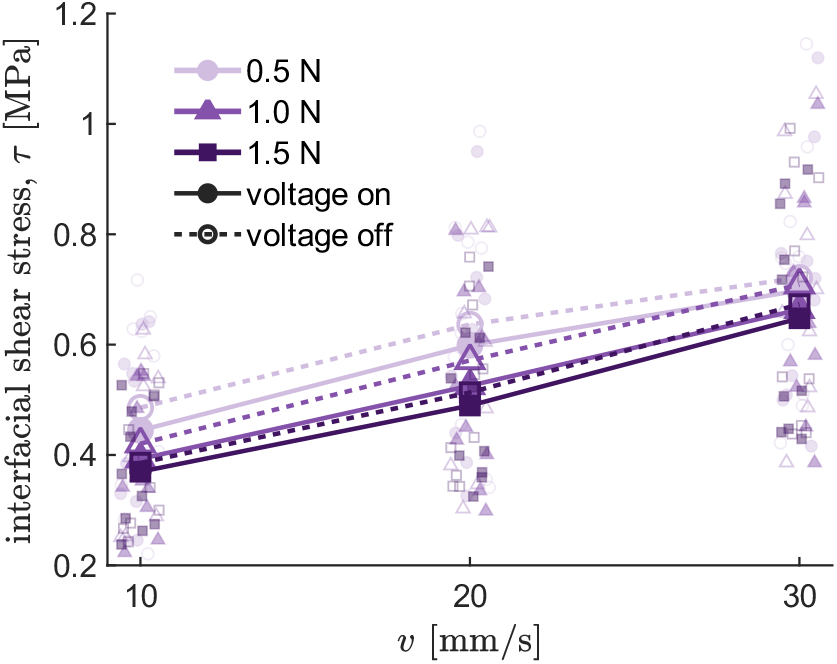
Interfacial shear stress (*τ*) across sliding speed (*v*), normal force (*F*_*n*_), and electrostatic actuation. Filled markers and solid lines indicate voltage-on conditions; open markers and dashed lines indicate voltage-off conditions. Small markers show participant-level values; large markers show means.

To quantify the trends in Fig. 2, we fitted a linear mixed-effects model of interfacial shear stress as a function of sliding speed, normal force, and their interaction, while accounting for repeated measurements across participants through a random intercept term. The model used the specification *Y* ~ 1+speed×force+(1|participant). The model revealed a significant positive effect of speed (*β* = 0.0109±0.00260 MPa per mm/s, *p* < 0.001) and a significant negative effect of force (*β* = −0.143 ± 0.0520 MPa/N, *p* < 0.05). In contrast, the interaction between speed and force was not significant (*p* = 0.283), indicating that there was no evidence that the effect of speed differed across the tested force range. A likelihood-ratio comparison showed that including a participant-level random intercept significantly improved model fit compared with a fixed-effects-only model (*p* < 0.001), indicating substantial inter-individual variability in the overall magnitude of interfacial shear stress. Consistent with this variability, participant-level mean interfacial shear stress, averaged across the nine speed–force conditions, ranged from 0.34 to 0.81 MPa, which falls within the range reported by Bochereau et al. [16].

To clarify the mechanical origin of the trends observed in Fig. 2, we analyzed the relative changes in real contact area and tangential force, the two quantities that define interfacial shear stress through *τ* = *F*_*t*_*/A* [9], with respect to reference operating points (Fig. 3). For the speed-dependent analysis, each participant’s values were normalized to the corresponding value at 10 mm/s within the same force level. For the force-dependent analysis, each participant’s values were normalized to the corresponding value at 0.5 N within the same speed condition. The resulting relative changes were calculated as Δ*X* = (*X/X*_*ref*_ − 1) × 100, where *X* denotes either real contact area or tangential force.

**Fig. 3:**
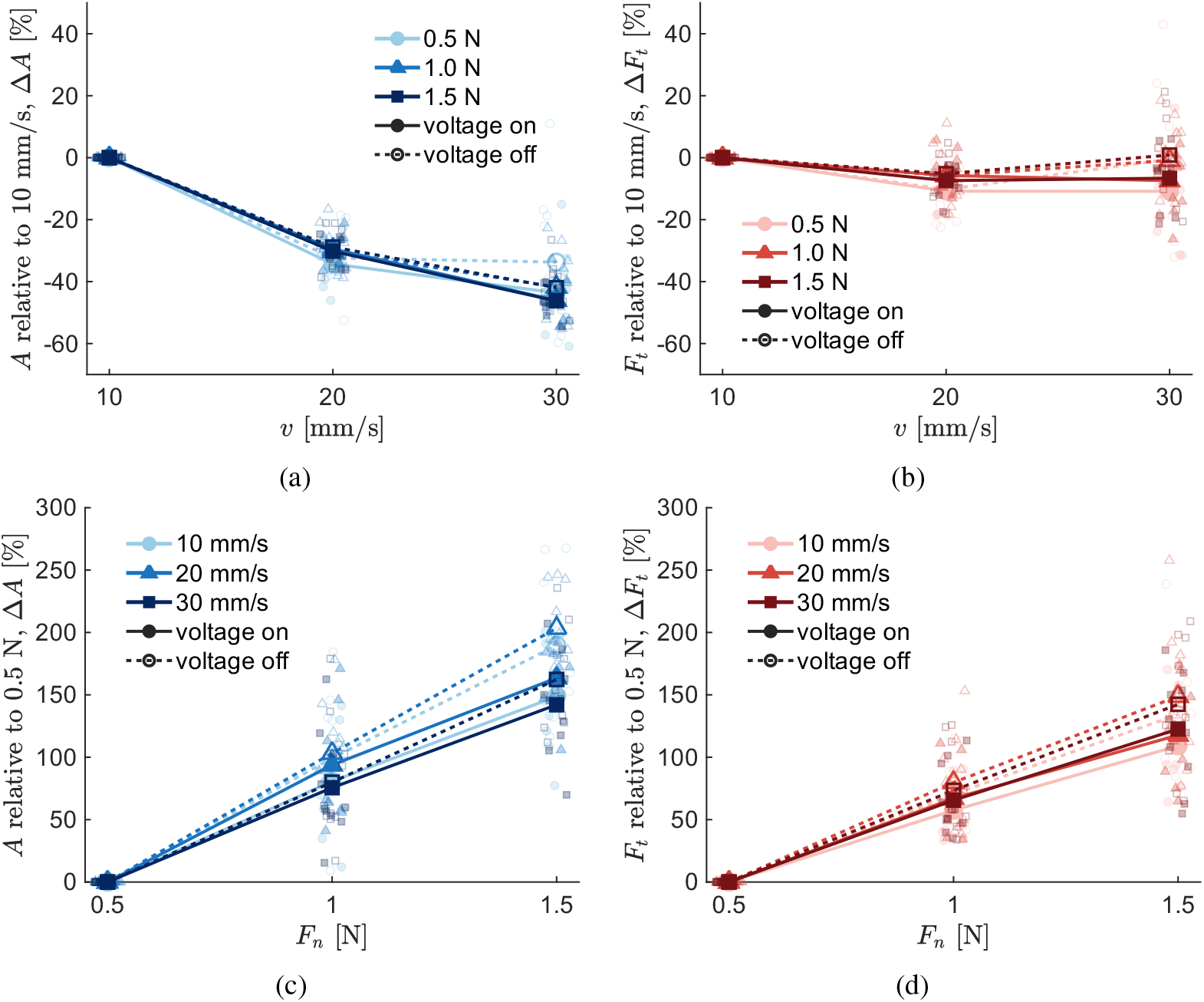
Relative changes in real contact area and tangential force across speed and normal force. Changes in (a) real contact area *A* and (b) tangential force *F*_*t*_ relative to 10 mm/s within each normal-force level. Changes in (c) *A* and (d) *F*_*t*_ relative to 0.5 N within each sliding-speed level. Filled markers and solid lines indicate voltage-on conditions; open markers and dashed lines indicate voltage-off conditions. Small markers show participant-level values; large markers and lines show condition means.

With increasing sliding speed, real contact area decreased more strongly than tangential force, indicating that the speed dependence of interfacial shear stress was primarily governed by the stronger variation in contact formation (Figs. 3a and 3b). By contrast, increasing normal force produced larger positive changes in real contact area than in tangential force, which explains why interfacial shear stress decreased with load despite the accompanying rise in tangential force (Figs. 3c and 3d). These results reveal that the observed interfacial shear stress trends arise from the competing contributions of contact growth and frictional force at the finger-surface interface.

As sliding speed increases at a constant normal force, real contact area decreases, which inherently raises the mean contact pressure (*p* = *F*_*n*_*/A*). The resulting increase in interfacial shear stress with sliding speed is therefore consistent with the pressure-dependency of shear stress reported in the literature for skin-surface interactions [13] and other materials [31], [32]. This reduction in contact area at higher velocities is also consistent with viscoelastic stiffening of the skin tissue; at higher deformation rates, the skin does not have sufficient time to fully conform to the surface asperities [15], [19], [21], [26], [33]. Furthermore, the rise in shear stress is consistent with the adhesive contribution to friction described by Schallamach’s theory [15], [34], where interfacial resistance is modulated by the rate of bond formation and breakage. Specifically, for rubber-like viscoelastic materials, this theoretical framework describes the adhesive contribution to shear stress as rising with sliding velocity up to a characteristic velocity associated with the timescale of interfacial bond formation and breakage, beyond which it declines [15]. For fingertip friction, previous work has shown that the effect of sliding velocity is strongly condition-dependent and influenced by moisture state, occlusion, and tribological configuration [18]. In the data of Pasumarty *et al*., the dry-fingerpad case appears to reach a friction maximum at approximately 100 mm/s, whereas this characteristic speed shifts to lower values under wetter conditions [18]. Therefore, within the tested velocity range of 10–30 mm/s, our data are consistent with the increasing part of a velocity-dependent interfacial response.

Regarding the effect of normal force, our statistical analysis revealed a significant, albeit relatively small, negative main effect: interfacial shear stress slightly decreased as the applied load increased. This reduction is broadly consistent with the macroscopic friction behavior of human skin, where the friction coefficient (*µ* = *F*_*t*_*/F*_*n*_) often decreases with increasing load on smooth counter surfaces [1], [5], [20], [35]. Although *µ* and *τ* (*τ* = *F*_*t*_*/A*) are distinct quantities, they can be directly linked through the mean contact pressure (*p* = *F*_*n*_*/A*), yielding the relationship *µ* = *τ/p*. For rough multiasperity contacts, several studies show that the real contact area increases approximately proportionally to the normal load [30], [36], [37]. Under this approximation, *p* would remain roughly constant with the load, so the well-documented increase in *µ* at lower normal forces would correspond to a higher *τ*. Our measurements are consistent with this interpretation: increasing the normal force increased both the tangential force and the optically resolved real contact area, but the contact area increased more strongly than the tangential force, so the ratio *τ* = *F*_*t*_*/A* decreased.

As in the voltage-off condition, interfacial shear stress increased with speed and decreased with force under electrostatic actuation (Fig. 2). In addition to these operating-condition effects, electrostatic actuation produced a modest but statistically significant reduction in the area-normalized shear stress. The participant-level mean ratio *τ*^*on*^*/τ*^*off*^, averaged across the nine speed–force conditions, was 0.949, corresponding to a mean reduction of 5.14% (95% CI: 2.09%–8.18%). A one-sample *t*-test against zero reduction confirmed that this decrease was significant (*t*(9) = 3.81, *p* = 0.0041), and the Wilcoxon signed-rank test yielded a consistent result (*p* = 0.0098). All condition-wise mean ratios were below unity, with reductions ranging from 3.10% to 7.75%. Participant-level changes ranged from a 10.17% decrease to a 5.49% increase, indicating inter-individual variability in the magnitude and direction of the voltage-induced shear-stress modulation. A mixed-effects model applied to the percentage reduction showed no significant effects of sliding speed, normal force, or their interaction, indicating that the reduction was broadly consistent across the tested sliding conditions.

The sliding-condition dependence of interfacial shear stress under electrostatic actuation can be interpreted through the same force–area balance, *τ* = *F*_*t*_*/A*, used for the voltage-off condition. The voltage-on state showed qualitatively similar speed- and force-dependent patterns: at higher sliding speeds, the relative reduction in real contact area was more pronounced than the corresponding change in tangential force, leading to a net increase in *τ* (Figs. 3a and 3b). In contrast, increasing normal force produced larger positive changes in real contact area than in tangential force (Figs. 3c and 3d), thereby explaining the decrease in interfacial shear stress despite the higher frictional load.

The reduction in *τ* caused by electrostatic actuation can be understood as an additional shift in this force–area balance. Electrostatic attraction can increase the real contact area, while the oscillatory normal forcing associated with actuation may weaken the effective tangential coupling at the interface. This may promote partial stick–slip or intermittent slip, limiting the increase in effective tangential force relative to the increase in real contact area [17]. As a result, the tangential-force increase under voltage-on can remain smaller than the accompanying increase in real contact area, producing a lower area-normalized shear stress. This interpretation is consistent with previous frequency-dependent analysis, where the vibration-dominated regime produced a reduction in interfacial shear stress under electrostatic actuation [17]. It is also consistent with mechanical vibration studies of finger–glass contact, which likewise attribute vibration-induced friction changes primarily to the tangential force term rather than to contact-area changes, although in that case the reported effect was a net decrease in tangential force rather than a limited increase [16].

We observed inter-individual variability in the reduction of *τ* caused by electrostatic actuation. One possible contributor is variation in skin hydration. Notably, the only participant showing an average increase in *τ* under electrostatic actuation also exhibited visible condensation, suggesting a more hydrated contact state [17], [38]. Several mechanisms could explain how hydration limits the shear stress reduction. First, moist fingers can exhibit larger baseline contact area and friction, leaving less room for additional contact-area modulation under electrostatic actuation [3], [17], [38]. Second, increased moisture may also introduce greater effective damping in the fingertip contact, limiting vibration-induced vertical displacement and reducing the modulation of tangential force and real contact area [17]. Third, finger moisture can alter the electrical coupling at the finger–touchscreen interface by reducing impedance and increasing conductivity, which may weaken the effective electrostatic modulation [17], [38].

The voltage-sweep experiment further supported this force– area interpretation (Fig. 4). Increasing voltage produced a progressive increase in both tangential force and real contact area ratios. However, the increase in real contact area was larger than the increase in tangential force, especially at higher voltages, so the area-normalized shear-stress ratio *τ*^*on*^*/τ*^*off*^ decreased with voltage. This trend is consistent with the voltage dependence of electrostatic attraction, which is expected to scale with *V* ^2^ according to parallel capacitance theory [5], [23], [27]: as electrostatic attraction increases, the normal oscillatory forcing at the fingertip–surface interface becomes stronger, potentially promoting partial stick–slip or intermittent slip that limits the relative increase in tangential force while allowing a larger increase in real contact area.

**Fig. 4:**
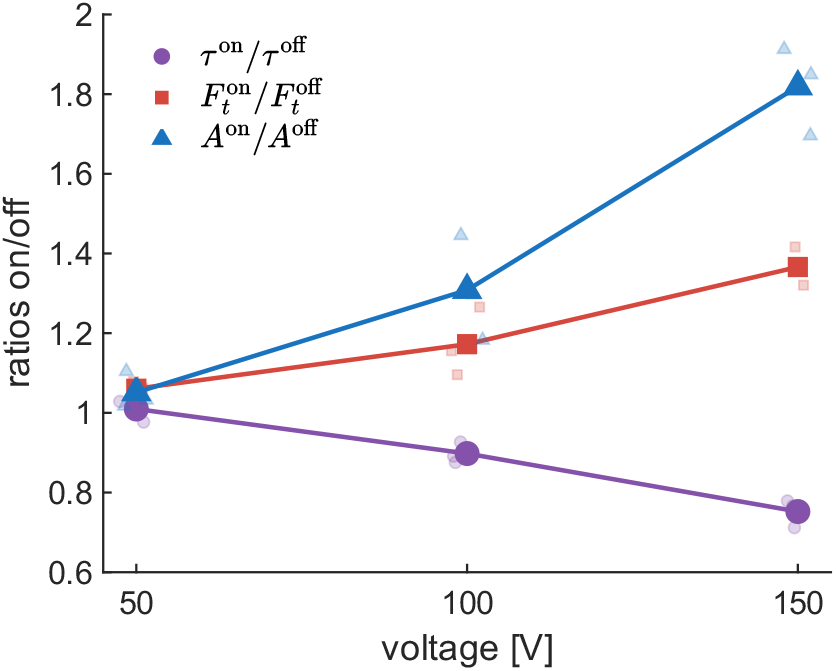
Voltage-dependent changes in interfacial shear stress, tangential force, and real contact area. Voltage-on/voltage-off ratios are shown for a voltage-sweep experiment performed at 20 mm/s and 1 N. Small transparent markers show individual repetitions; large markers show mean values.

## IV. Conclusion

This study experimentally showed that the mean interfacial shear stress of a sliding fingertip is condition-dependent. By combining tangential force and optically resolved real contact area measurements, we found that interfacial shear stress increased with sliding speed and decreased with normal force. These trends were explained by the force–area balance: speed reduced real contact area more strongly than tangential force, whereas normal force increased real contact area more strongly than tangential force. Electrostatic actuation produced a modest reduction in interfacial shear stress; real contact area increased more strongly than tangential force with actuation amplitude. The magnitude of these effects varied across participants, potentially reflecting differences in skin hydration and other individual contact properties. These results clarify how changes in real contact area and interfacial shear stress jointly shape fingertip–surface friction.

